# Estimating Actual Striking Forces Using Attenuation Properties of Taekwondo Protectors

**DOI:** 10.1101/2025.07.13.663825

**Authors:** Minho Chae, Jonghak Hwang, Woosup Han, Sihyun Ryu, Sangkyoon Park, Sukyung Park

**Affiliations:** Department of Mechanical Engineering, Korea Advanced Institute of Science and Technology (KAIST), Yuseong-gu, Daejeon, Republic of Korea; Research Department of Sport Industry, Korea Institute of Sport Science (KISS), Songpa-gu, Seoul, Republic of Korea; Technology Innovation Team, Korea Sports Promotion Foundation, Kiheung-gu, Yongin-si, Gyeonggi-do, Republic of Korea; Motion Innovation Center, Korea National Sport University, Songpa-gu, Seoul, Republic of Korea

## Abstract

**Background:** Protectors attenuate impact forces significantly, however, their quantitative attenuation characteristics under varying conditions remain poorly understood. Accurate measurement of striking forces is essential for evaluation. However, direct measurement of athletes’ striking forces presents significant practical and safety challenges, necessitating the development of indirect estimation approaches.

**Method:** A controllable pendulum-based impact testing apparatus was developed to evaluate force attenuation characteristics of Taekwondo body protectors. The system delivered repeatable impacts across varying magnitudes and contact durations to a mannequin equipped with a protector. Simultaneous measurements of input and transmitted forces through the protector enabled direct quantification of attenuation ratios under controlled conditions. To demonstrate practical application, five Taekwondo athletes performed standardized kicks on the instrumented protector setup, allowing estimation of their actual striking forces using the derived attenuation factor.

**Results:** Force attenuation ratios demonstrated high consistency across impact magnitudes ranging from 300 to 5800 Newtons, with impact duration variations showing minimal influence on attenuation performance. Linear attenuation relationships were established between input and transmitted forces (*R*^2^*≥* 0.987), enabling derivation of a predictive attenuation factor. Validation testing with the pendulum system showed that the estimated input force trajectories achieved an nRMSE of 3.5%. Averaged across the entire force range, the MAPE was 4.7% for maximum force, 1.4% for impact duration, and 3.5% for impulse. Applying the attenuation factor to standardized kicks performed by five Taekwondo athletes yielded estimated maximum striking forces ranging from approximately 1300 to 1800 Newtons, demonstrating the method’s capability to quantify actual forces that would otherwise be difficult to measure directly.

**Discussion:** This study establishes the first quantitative characterization of impact attenuation in Taekwondo body protectors, providing a validated attenuation factor for estimating striking forces from transmitted forces. These findings demonstrate a practical methodology for quantifying striking forces that are otherwise difficult to measure directly.

**Author summary:** 

## Introduction

In recent years, global interest and participation in combat sports have continued to rise [1]. In disciplines such as Taekwondo, precise measurement of an athlete’s striking force is indispensable for performance analysis, training optimization, injury prevention, and related objectives. To measure these forces while safeguarding athletes, the force sensors are routinely covered with a protector or cushioning material. However, since the protector absorbs and attenuates part of the applied force, considerable discrepancies arise between the actual striking force delivered by the athlete and the force measured by the sensors. This attenuation effect continues to pose significant challenges for accurately quantifying striking forces while wearing protective gear.

Previous studies have employed diverse systems including water-filled heavy bag systems [2, 3], force plate systems [4–6], strain gauge-based systems [7–9], accelerometer-based systems [10–12], load cell systems [13–15]. Nevertheless, large discrepancies—often exceeding an order of magnitude—have been reported between maximum force values depending on the measurement system used, highlighting ongoing limitations in measurement consistency and reliability. In many studies, the protective padding or damping material mounted on top of the force sensor differs in thickness and composition, causing the cushioning effect to vary across experimental protocols. This variability makes it difficult to measure striking forces accurately and consistently, or to compare results between studies. Alternatively, several studies have measured impact forces using methods that entirely eliminate the cushioning effect [16–18], most commonly by estimating impact forces from pre-impact velocity rather than by employing force transducers. However, such indirect approaches, which estimate force from pre-impact velocity or acceleration, can only provide total impulse and fail to capture the force–time profile, such as maximum force or contact duration. They also neglect the complex multi-joint dynamics involved in actual striking movements, limiting their accuracy. Some of these studies attempted to directly measure impact forces by attaching flexible pressure sensors to the foot or striking body part, aiming to avoid injury risk [30–32]. However, such sensors typically estimate the total contact force based on localized pressure readings, which may not accurately represent the distributed force profile across the entire contact surface, especially under dynamic impact conditions. Conversely, several studies have employed striking machines to evaluate the damping effect of protective equipment such as boxing gloves; however, most of these works attached sensors to only one side of the impact apparatus (either the striker or the target) and inferred the forces and impulses acting on the opposite side indirectly [19–25]. To the best of our knowledge, no previous study has directly quantified the attenuation of force profiles through protectors by simultaneously mounting sensors on both the striking and impacted sides. The present work addresses this gap by instrumenting the inner (impacted) and outer (striking) surfaces of the protector with force transducers, thereby enabling accurate estimation of the true force profile and systematic quantification of the protector’s force attenuation characteristics. In this study, we investigated how input striking forces are attenuated by Taekwondo protectors. To directly quantify impact magnitude and contact dynamics through force transmission, we developed a pendulum-based impact apparatus capable of replicating a range of striking scenarios with coupled variations in force magnitude and contact duration. By instrumenting force transducers on both the striking and impacted surfaces, we directly measured input and transmitted force profiles. From these data, we derived consistent attenuation factors that characterize the force attenuation properties of the protector. Finally, we applied these factors to transmitted force measurements obtained from actual Taekwondo athletes performing kicks, thereby enabling estimation of the original striking forces that would otherwise be difficult to measure directly.

## Materials and methods

The experimental procedure consisted of three main stages: (1) development of a pendulum-based impact apparatus, (2) derivation of attenuation factors through controlled impact testing, and (3) estimation of athletes’ striking forces based on kicking experiments.

### Impact System Development

#### Impulse–Momentum–Based Design Principles

The following impulse–momentum relationships were used to design and analyze the pendulum-based impact system. When the pendulum is assumed to be stationary before impact, the impulse *J* is defined as:

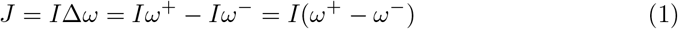

where *I* is the pendulum’s moment of inertia, and *ω*_+_ and *ω*_*−*_ are the angular velocities immediately after and before impact, respectively.

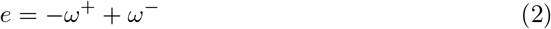

Using this definition, the impulse can be expressed as a function of the angular velocity at impact

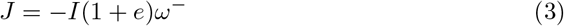

Hence, if the coefficient of restitution *e* is known and the impact angular velocity of the pendulum is measured, the impulse can be calculated. Assuming pivot friction and other energy losses of the pendulum are negligible, the work–energy principle allows the angular velocities to be determined from the initial drop angle *θ*_*i*_ and the rebound angle *θ*_*f*_

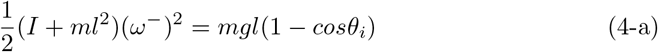

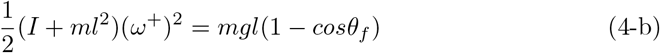

where *m* is the mass of the pendulum, *g* is the gravitational acceleration, and *l* is the effective length of the pendulum to the center of mass. From Eq (1),(2),(4-a) and (4-b), the impulse can be written as

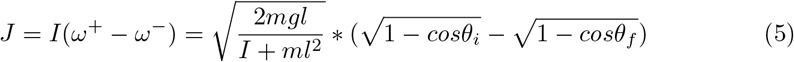

Based on this relationship, the pendulum’s mass, effective length, and initial drop angle were chosen to achieve the target impulse range in the design.

#### Impactor unit

To generate simple and repeatable impact forces, an impactor unit based on the free-fall pendulum principle was developed (Fig 1A, 1B). The system parameters were determined to achieve impulse values in the range of approximately 15–50 Ns, based on previously reported maximum forces and contact durations observed in human strikes (Table 1). Because impulse is governed by physical design parameters such as mass, moment of inertia, and release angle, the system enables stable and reproducible impact generation. The vertical pendulum was designed with adjustable mass of 10–15 kg, and a steel frame of 1.04 meters in length to ensure structural durability under repeated high-intensity impacts. The striking head, which contacts the target, was fabricated from aluminum, with a mass of 3.5 kg and a length of 0.385 meters (Fig 1B). The pendulum system allows fine manual adjustments of both horizontal and vertical alignment. A rotary encoder was integrated for real-time measurement of the release angle. The release and braking mechanism employs a powder clutch system to minimize structural vibration and to maintain consistent initial conditions throughout repeated trials.

**Table 1.** Summary of impact characteristics from previous and current studies on human striking actions. For each study, the type of measurement system and the corresponding striking action(s) are listed. When multiple actions were assessed within a study, they are separated by a tilde (*∼*). Summary data are reported as mean (SD)

| Study | Measurement | Action | Maximum Force (N) | Impact Duration (s) | Impulse (Ns) |
| --- | --- | --- | --- | --- | --- |
| Pieter and Pieter (1995) [2] | Water-filled heavy bag, built-in sensor | Taekwondo kicks and punch | 341 (44)<br>~ 661 (52) | – | – |
| Girodet et al. (2005) [27] | Force sensor | Karate punch | 1745 | 0.015 | 13.7 |
| Pedzich et al. (2006) [28] | Force plate | Taekwondo kicks | 7751 (2570)<br>~ 8569 (2381) | 0.016 (0.002)<br>~ 0.018 (0.004) | 60 (24)<br>~ 84 (30) |
| Gulledge & Dapena (2007) [29] | Force plate | Karate punches | 790 (130)<br>~ 1450 (290) | 0.022 (0.004)<br>~ 0.032 (0.008) | 37.7 (14.6)<br>~ 43.2 (15.3) |
| Falco et al. (2009) [30] | Film sensor | Taekwondo kick | 1304 (608)<br>~ 2089 (634) | – | – |
| Sullivan et al. (2009) [31] | Accelerometer | Taekwondo kick | 5419 (659)<br>~ 6400 (898) | – | – |
| Estevan et al. (2011) [32] | Film sensor | Taekwondo kick | 1203 (154)<br>~ 1829 (161) | 0.022 (0.001)<br>~ 0.029 (0.002) | – |
| Neto et al. (2012) [13] | Load cell | Palm strike | 957 (436)<br>~ 1884 (900) | – | – |
| Buško et al. (2014) [8] | Dynamometer | Boxing punch | 1634 (674)<br>~ 2899 (457) | – | – |
| Wasik & Nowak (2015) [17] | 3D imaging | Taekwondo punch | 1308 (173) | – | 0.170 (0.007) |
| Svoboda et al. (2016) [33] | Dynamometer | Boxing punch | 1970 ~ 2292 | 0.0109 ~ 0.0129 | – |
| Buško et al. (2016) [11] | Accelerometer | Boxing punch<br>Taekwondo kick | 848 (248)<br>~ 3514 (1190) | – | – |
| de Souza et al. (2017) [34] | Strain gauge | Karate punch | 1812 | – | – |
| Gavagan& Sayers (2017) [7] | Strain gauge | Taekwondo<br>Karate<br>Muay Thai kick | 1211 (219)<br>~ 1547 (530) | – | 30.9 (6.9)<br>~ 38.8 (15.3) |
| Lenetsky et al. (2018) [12] | Accelerometer | Punch | 1411 (365)<br>~ 4749 (107) | – | 26 (7)<br>~ 75 (8) |
| Jovanovski & Stappenbelt (2020) [14] | Load cell | Boxing punch | 758 ~ 3845 | – | – |
| Khasanshin & Aleksey (2021) [35] | Air Pressure | Boxing Punch | 1006 (198) | – | – |
| Beranek et al. (2022) [36] | Force plate | Punches<br>Elbow strike | 3158 (1217)<br>~ 6460 (1612) | – | 8.4 ~ 19.6 |
| Finlay et al. (2023) [37] | Force plate | Boxing punch | 1645 (537)<br>~ 2624 (581) | – | – |
| **Current study (estimated input)** | **Load cell** | **Taekwondo kick** | **1491.16 (238.28)** | **0.026 (0.002)** | **20.28 (4.77)** |

**Fig 1.**
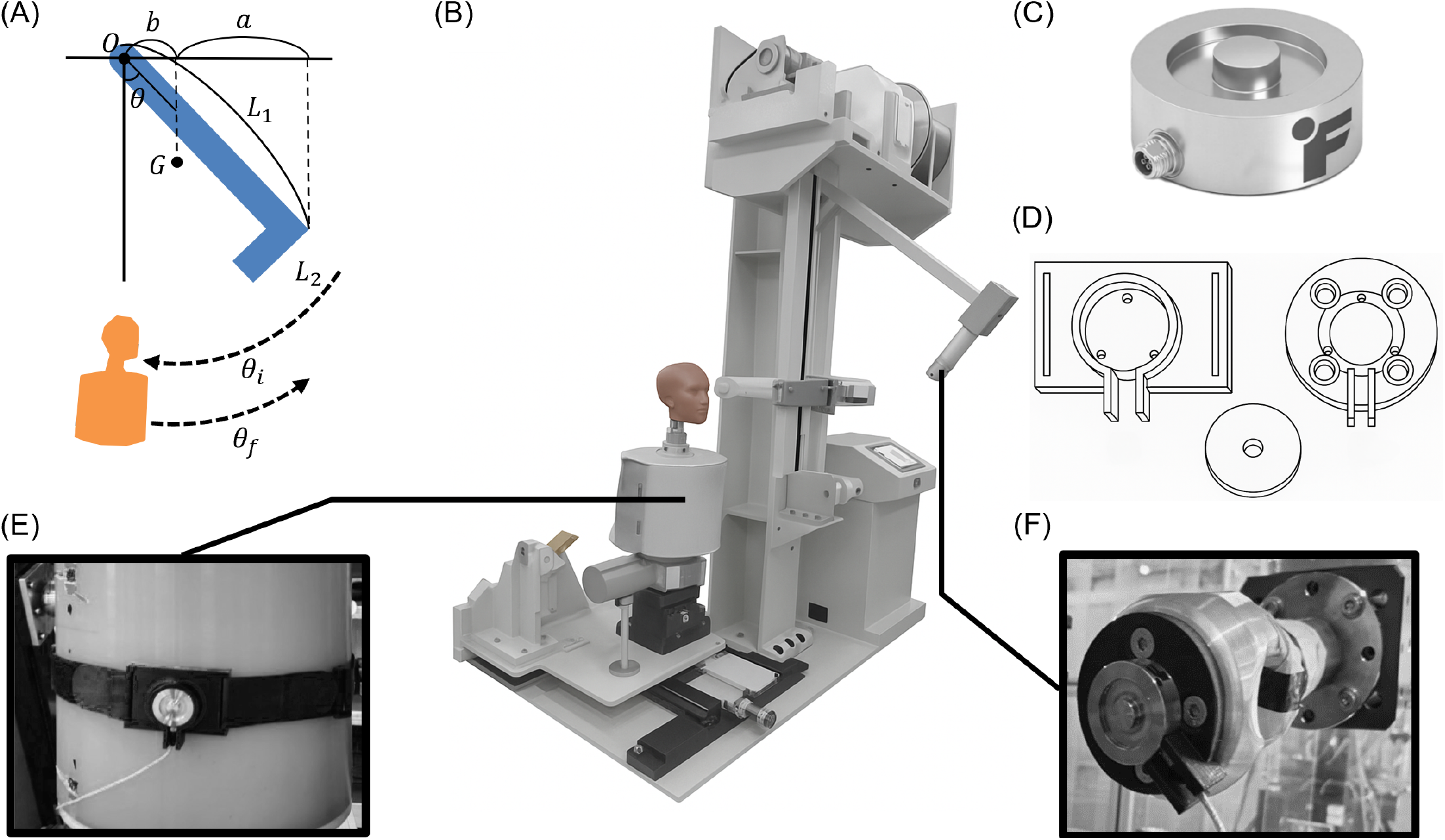
Experimental apparatus for measuring force attenuation in Taekwondo body protectors. (A) Design schematic of the pendulum-based impact apparatus, illustrating the swing trajectory and release angle (*θ*). (B) Overall view of the experimental setup, comprising a pendulum impactor, a mannequin outfitted with a Taekwondo body protector, and a force measurement system. (C) Load cell used for force measurement. (D) A custom-designed load cell adapter cap and mounting. (E) Transmitted force measurement using a load cell positioned behind the protector on the mannequin surface. (F) Input force measurement using a load cell mounted at the pendulum head.

#### Target unit

The target unit consisted of a cylindrical aluminum mannequin (6061 alloy) designed to approximate the geometry of a human upper torso (Fig 1B). The mannequin was mounted on a pivot to allow free rotation, simulating the backward displacement of the human body upon receiving a kick. After each impact, an automated motorized return system rapidly reset the target to its initial position, ensuring both efficiency and consistency across repeated trials.

#### Measurement Unit

Impact forces were measured using an LLB 450 load cell (FUTEK, USA; maximum load: 22,241 N, natural frequency: 34 kHz, material: 17-4PH) (Fig 1C). A custom-designed adapter cap was installed on the load cell to ensure consistent contact with the protector’s curved inner surface and minimize misalignment. Preliminary tests using an instrumented hammer confirmed that the cap introduced no significant errors under moderate loading (Fig 1D). Prior to the main experiments, quasi-static and low-speed impact tests using an instrumented hammer confirmed that the adapter cap did not introduce significant measurement discrepancies under moderate loading conditions. Sensor signals were amplified using an ST-AM100 amplifier (Sensetech, UK) and recorded via a USB-6211 DAQ (National Instruments, USA). Data acquisition and control were performed using LabVIEW software. Input impact forces were measured using the load cell mounted on the pendulum head (Fig 1F), while transmitted forces were simultaneously measured by the load cell mounted behind the protector on the mannequin (Fig 1E). This dual-sensing configuration allowed direct quantification of the protector’s impact attenuation performance. Here, the ‘input force’ (also referred to as ‘striking force’) is defined as the actual force exerted by the striker onto the protector surface, and the ‘transmitted force’ as the attenuated force measured behind the protector. For clarity, ‘input force’ and ‘striking force’ are used interchangeably.

### Derivation of Attenuation Factors

To evaluate the impact attenuation characteristics of the body protector, repeated impact tests were conducted across a range of release angles, varying the pendulum’s initial angle from 20° to 80° in 10° increments. For each angle, five impact trials were performed in both increasing and decreasing angle sequences, resulting in five repetitions per condition. Force data collected from both the pendulum head and the load cell mounted behind the protector on the mannequin were sampled at 1000 Hz. To reduce drift and high-frequency noise, the signals were smoothed using a fourth-order Butterworth bandpass filter with a passband of 0.5–350 Hz.

Three metrics were defined to quantitatively assess the protector’s attenuation performance: Maximum Force, Impact Duration, and Impulse. Maximum Force was defined as the peak force recorded during each impact. Impact Duration was determined by identifying the time interval between the first and second instances when the force exceeded 2% of the maximum value. Impulse was calculated by integrating the force-time curve between these two time points.

To examine the linearity between input and transmitted forces, linear regression analyses were performed for each metric. To confirm linearity, the coefficient of determination (*R*^2^) was required to be 0.95 or greater. From the resulting linear relationships between the input forces (measured at the pendulum head) and transmitted forces (measured at the back of the protector), attenuation factors were derived for each impact parameter.

### Athlete Kicking Experiments

To estimate the actual striking forces of athletes based on the previously derived attenuation factors, additional kicking experiments were conducted. Five elite male Taekwondo athletes from Chungnam National University (height: 180.76 ± 4.27 cm, mass: 70.28 ± 3.56 kg, age: 20.6 ± 0.89 years) participated in the study. All participants were recruited on 29 May 2024 and were adult collegiate elite Taekwondo athletes. After receiving a full explanation of the study objectives and procedures, each participant provided written informed consent, and the study was approved by the KAIST Institutional Review Board (approval number: KH2024-089).

Following a 20-minute warm-up, each athlete performed 20 roundhouse kicks, the signature kicking technique in Taekwondo [26], against a sandbag equipped with the body protector used in previous tests. Athletes were instructed to perform kicks as they would in actual competition scenarios. The impact forces generated by the kicks were measured using an LLB 400 load cell (FUTEK, USA; maximum load: 10,020 N, natural frequency: 30.7 kHz, material: 17-4PH) mounted inside the sandbag and sampled at 1000 Hz. The load cell height was individually adjusted for each participant based on their body height and leg length. To ensure accurate targeting, the impact zone of the load cell was visually marked on the protector surface, and ink was applied to the participants’ feet to verify contact location after each trial. Among the total 100 kicks performed, 50 trials (11, 18, 6, 3, and 12 trials per athlete, respectively) in which the ink-marked region confirmed direct contact with the effective sensing area of the load cell were included in the final analysis. Raw force data were analyzed without applying additional filtering to minimize the distortion of force profile.

## Results

The impact-force profiles generated by the developed pendulum-based impact apparatus covered the range of force magnitudes and impact durations reported in previous studies on human Taekwondo kicks (Fig 2A, Table 1). By varying the pendulum’s free-fall drop height, our impact apparatus replicated 80–90% of the spectrum of impact forces (0.34–8.5 kN) and impact durations (10–32 ms) reported in previous studies. With increasing release angles, corresponding to higher collision velocities, maximum impact forces increased while impact durations decreased (Fig 2B, 2C). At lower angles, force profiles exhibited a triangular-like shape, whereas at higher angles, a short plateau was followed by a steep rise and rapid decay of the impact force.

**Fig 2.**
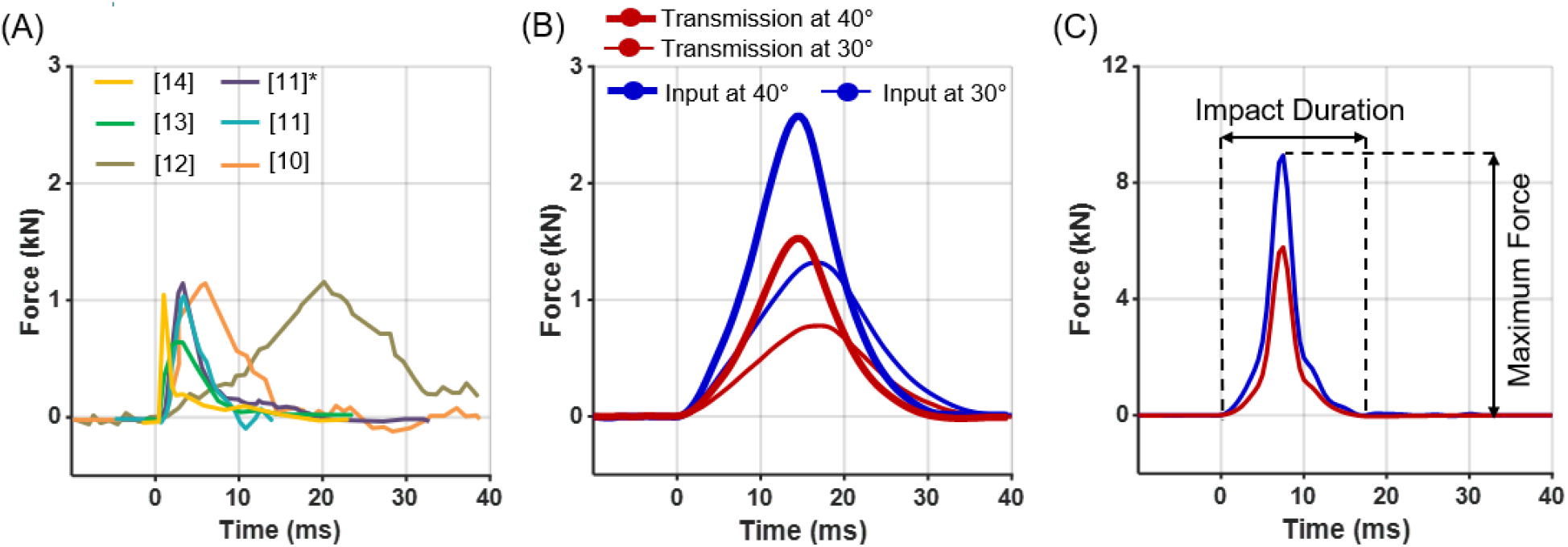
Profiles of impact force. (A) Reference force profiles of various strikes reported in previous studies, including Taekwondo kicks. (* Other strike types examined in the same study.) (B) Force profiles obtained from our pendulum impact apparatus at 30° and 40° release angles (thick solid lines). Input forces (blue) and transmitted forces through the body protector (red) are shown. (C) Force profiles from 80° free fall impacts, illustrating the definitions of maximum force and impact duration.

The impulse analysis of the pendulum-based impact apparatus demonstrated high consistency with the theoretical values used during system design (Fig 3). The pendulum lost energy during impact and rebounded to approximately 60% of its initial drop angle (Fig 3). The coefficient of restitution, e, calculated using the impulse-momentum relationship (Eq 1), showed a decreasing trend with increasing release angle (i.e., impact velocity) (Fig 3B). Although e is often assumed to be constant, the observed decrease at higher impact velocities likely reflects increased deformation and energy dissipation between the pendulum and the protector during high-speed collisions. The impulse exerted on the mannequin, calculated from the change in angular momentum of the pendulum, increased proportionally with impact velocity (Fig 3B). Compared to the impulse estimated under a constant e = 0.6 assumption, the experimentally observed reduction in e resulted in slightly higher impulse values (Fig 3C).

**Fig 3.**
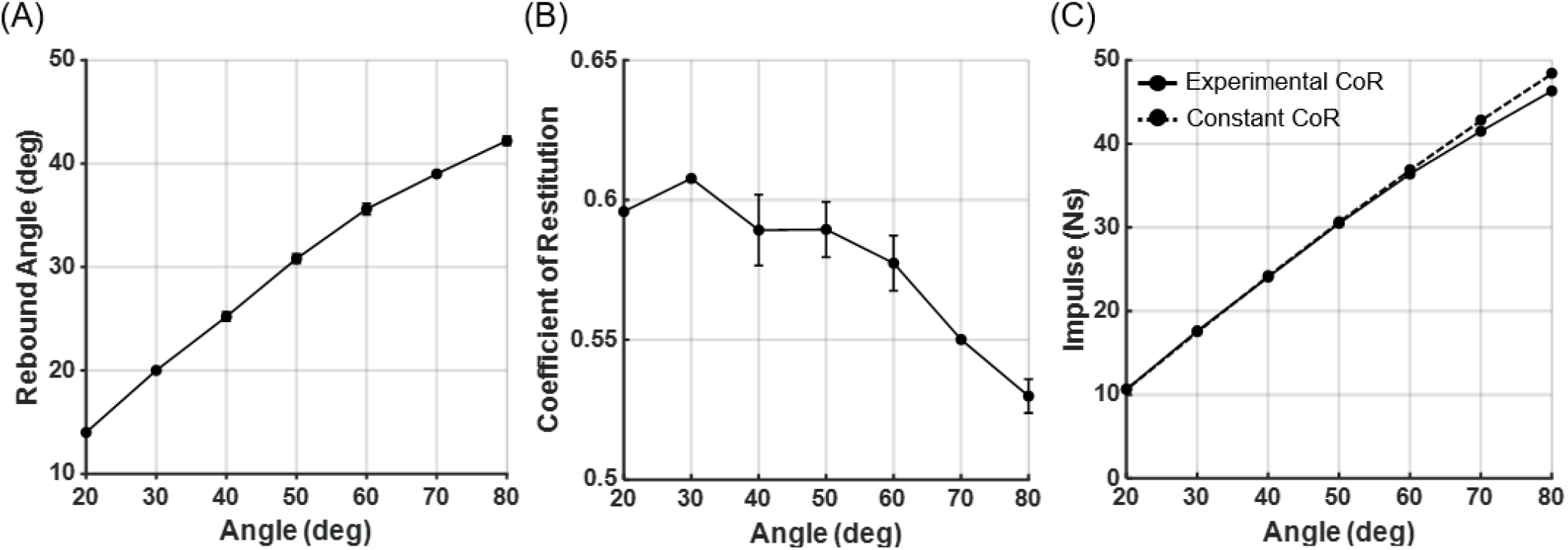
Performance evaluation of the impact apparatus. (A) Rebound angle of the pendulum after impact. (B) Experimentally calculated coefficient of restitution, e. (C) Corresponding estimation of impulse applied to the mannequin based on the pendulum motion. The dashed line in (C) indicates impulse estimation assuming a constant e = 0.6. Black dots represent the mean of five trials, and error bars indicate standard deviation

Both input forces and transmitted forces through the protector exhibited similar trends with increasing pendulum release angle (Fig 4). In both cases, maximum impact force increased proportionally with release angle (i.e., impact velocity), while impact duration decreased accordingly. By varying drop heights, the developed system generated combinations of impact force and contact duration that encompass most of the impact conditions reported for Taekwondo kicks in previous studies.

**Fig 4.**
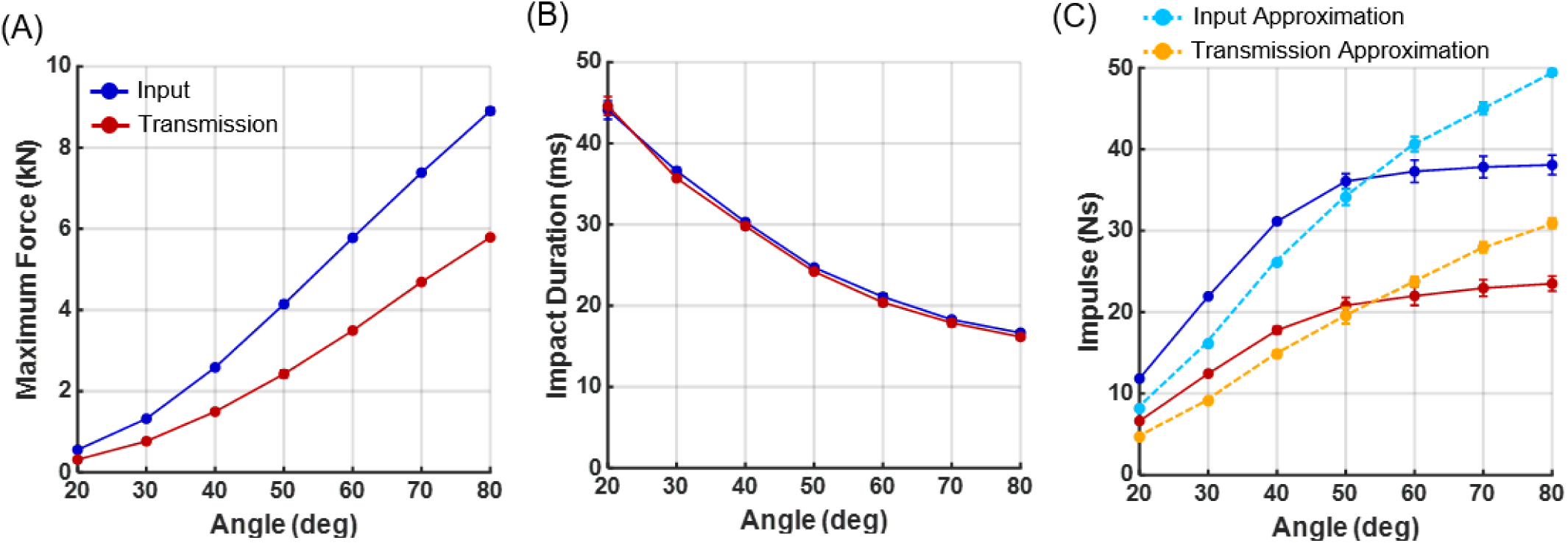
Input and transmitted impact-force metrics as a function of pendulum release angle. Experimental results of input (blue) and transmitted (red) forces. (A) Maximum impact force, (B) impact duration, and (C) impulse as a function of pendulum release angle. Approximated impulses, calculated as the product of maximum force and impact duration, are also shown for input (light blue) and transmitted (orange) forces. Each data point represents the mean of five repeated trials, with error bars indicating standard deviation. Overall, the very small error bars demonstrate the high repeatability of the test protocol.

However, with increasing impact velocity, the measured impulse values showed a tendency to plateau, which contrasted with the predicted impulse estimated from the difference in pendulum height before and after impact (Fig 3). Impulse is calculated as the integral of the force-time trajectory. When the force profile is approximated as a triangular waveform, impulse can be simply estimated as the product of maximum force and impact duration. This approximation yields a linear increase in impulse with release angle (impact velocity), which closely matches the impulse estimated from the pendulum’s rebound height (Fig 3).The observed plateau of measured impulse at higher velocities can be attributed to the steep force profiles observed in (Fig 2), where the brief duration of elevated force reduces the total integrated impulse. These results suggest that the impact forces measured by the load cell attached to the pendulum head may underestimate the true impulse during high-speed impacts. This discrepancy may arise from several experimental factors, including limitations in the dynamic response of the force transducer, signal filtering, and compliance in the sensor mounting structure. More detailed explanation of these factors is provided in the discussion section.

Across a broad range of impact conditions, the transmitted forces measured through the body protector demonstrated highly consistent relationships with the corresponding input forces (Fig 5). Maximum impact force and impulse were attenuated by approximately 60% through the protector, whereas impact duration was almost identical between input and transmitted forces. The consistency and repeatability of these relationships enable the derivation of attenuation factors that can estimate the input force profile based on transmitted force measurements. The derived attenuation factors, the ratio of the transmitted force to the input force, were approximately 0.63 for maximum force, 0.99 for impact duration, and 0.59 for impulse.

**Fig 5.**
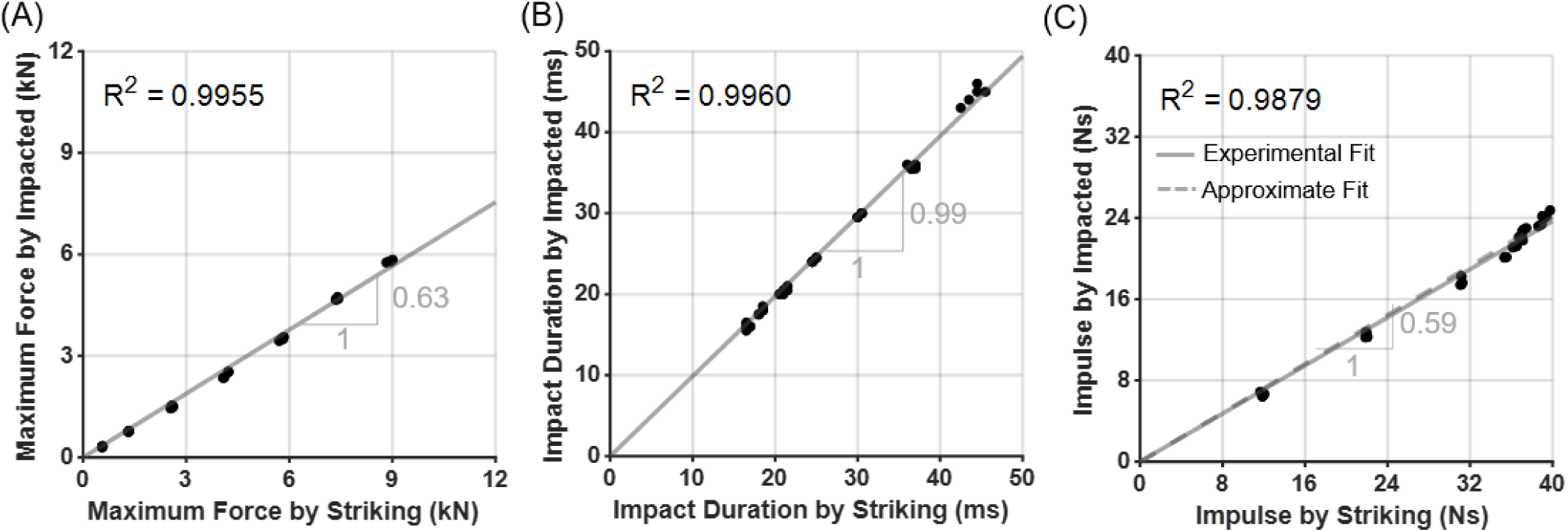
Relationships between input and transmitted impact parameters. The ratios between transmitted and input (A) maximum impact force, (B) impact duration, and (C) impulse are shown. Black dots represent individual test trials, and grey lines indicate linear regression fits.

The estimated input forces, derived from the transmitted forces using the previously established attenuation factors, reproduced the input force profiles well (Fig 6A). The full force–time trajectory showed a mean nRMSE (normalized Root Mean Squared Error) of 3.5%, while the impact characteristics exhibited MAPE (Mean Absolute Percentage Error) within 15% for maximum force, 6% for impact duration, and 10% for impulse. Higher input force magnitudes were associated with improved estimation accuracy for maximum force, while no consistent trend was observed for impact duration and impulse. These estimation errors are primarily governed by the attenuation factors determined in Fig 5. For instance, applying a regression weighting that better reflects the 0–3 kN range, which corresponds to the typical impact forces reported for Taekwondo kicks, could enhance estimation accuracy within this lower force range; under this weighting, the mean maximum force error within the band falls to just 2.2%.

**Fig 6.**
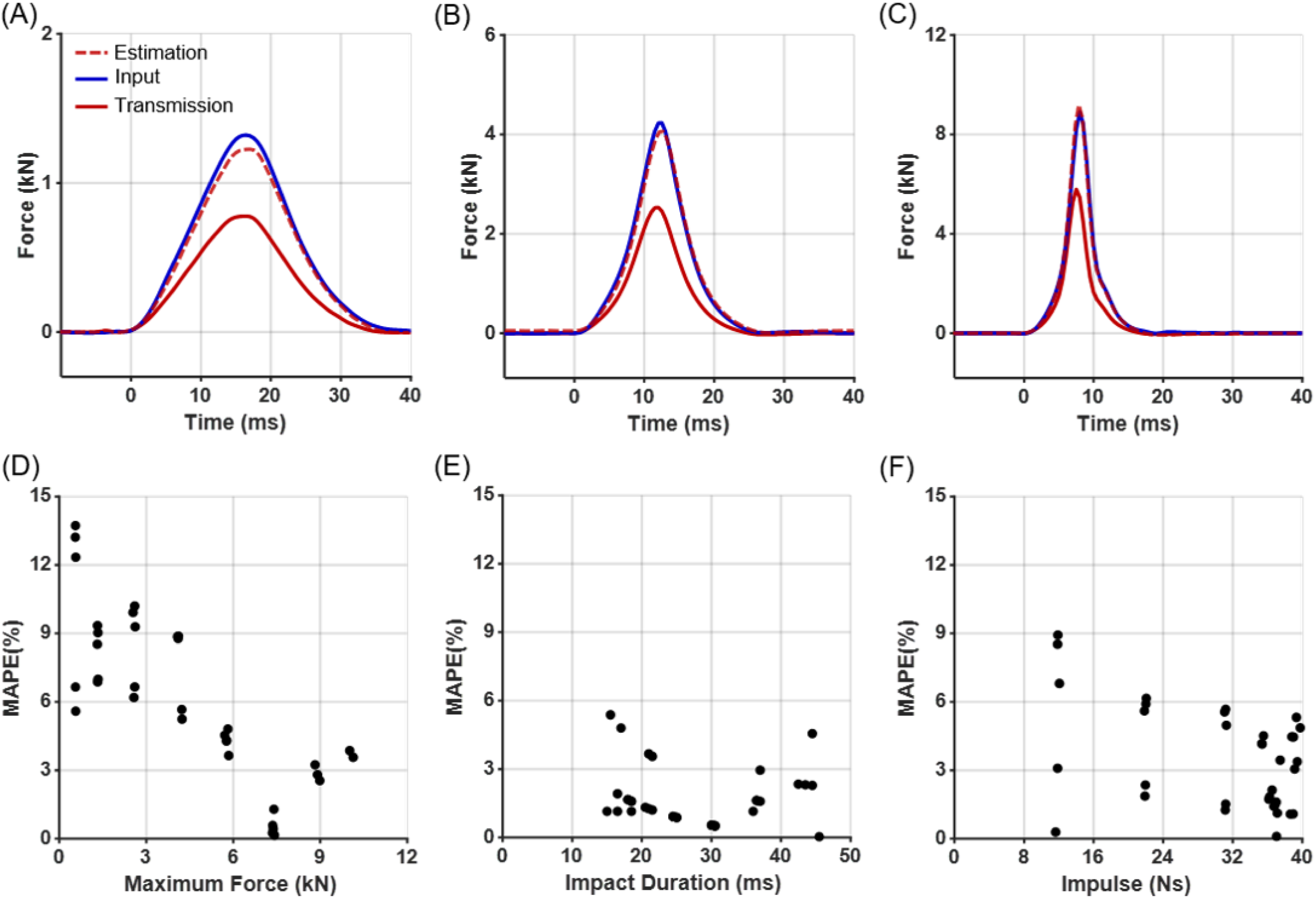
Validation of force estimation based on transmitted forces. (A-C) Examples of force profiles for input (blue), transmitted (red), and estimated input forces (red dashed) derived from transmitted forces using the attenuation factors. (D-F) Corresponding estimation errors for (D) maximum force, (E) impact duration, and (F) impulse, expressed as mean absolute percentage error (MAPE).

We applied the derived attenuation factors to athlete kick data to estimate the original striking forces (Fig 7). The kick force profiles were broadly categorized into three types: double-peaked profiles with a dominant first peak (Fig 7A), double-peaked profiles with a dominant second peak (Fig 7B), and single-peak profiles (Fig 7C). Double-peaked profiles accounted for the majority of kicks (92%), with the first peak being larger in 46% of cases and the second peak larger in 46% of cases. The maximum transmitted forces during kicks were approximately 0.939*±* 0.15 kN, with corresponding impact durations of 25.41*±* 2.33 ms and impulses of 12.02*±* 2.83 Ns. These ranges correspond to the low-intensity region of the impact apparatus (release angle 20-30°) for maximum force, the moderate-intensity region (release angle 40-50°) for impact duration, and the low-intensity region (release angle 30°) for impulse, respectively. Using the mean impact-force profile, the kick’s peak impact force was estimated to be approximately 1.5 kN, with a corresponding impact duration of approximately 25 ms and an impulse of approximately 20 Ns.

**Fig 7.**
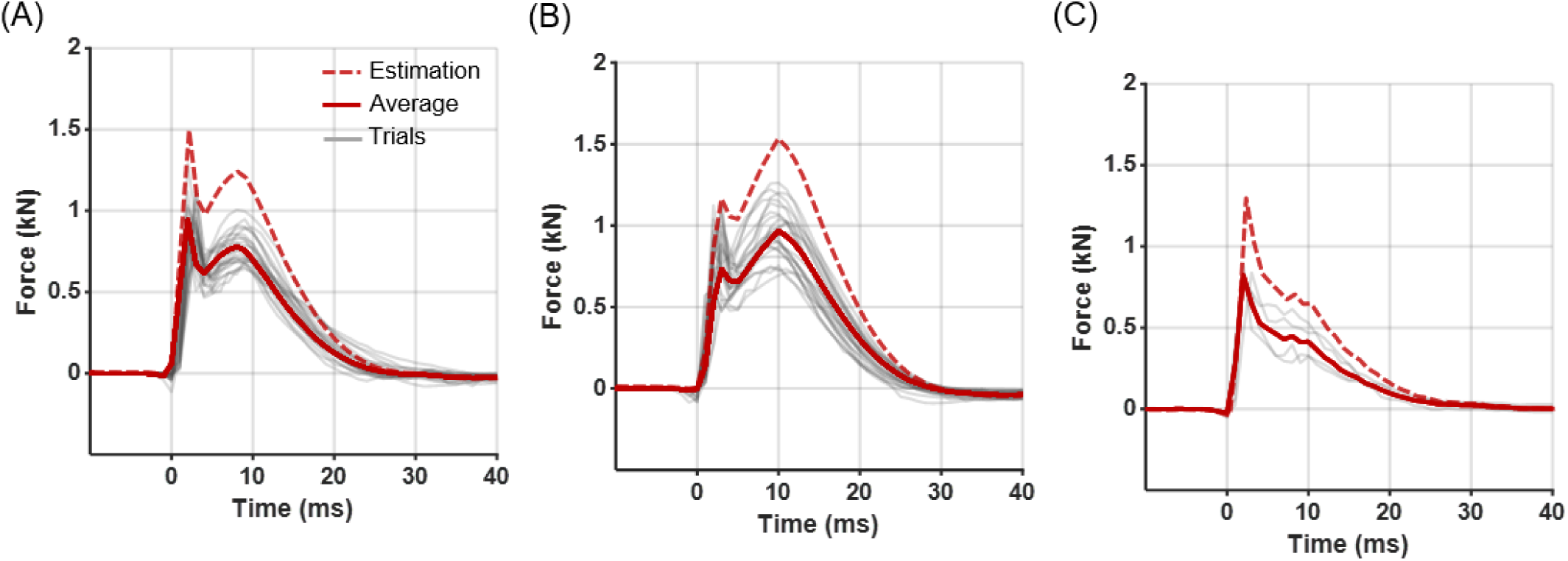
Measured and estimated Taekwondo kicks. The measured kick force profiles were categorized into (A) double-peaked profiles with a dominant first peak, (B) double-peaked profiles with a dominant second peak, and (C) single-peak profiles. Thin grey lines represent individual trials, red solid lines indicate the averaged transmitted force profiles, and red dashed lines show the estimated input force profiles reconstructed using the attenuation factors.

## Discussion

### Key Findings

This study presents, to our knowledge, the first attempt to quantitatively evaluate the impact attenuation performance of Taekwondo body protectors using a free-fall pendulum system under repeatable dynamic impact conditions, and to extend this evaluation to estimate actual striking forces in athletes. The experiments demonstrated that both maximum force and impulse transmitted through the protector were attenuated by approximately 60%, while the impact duration remained largely unchanged. A highly linear relationship was consistently observed between input and transmitted forces across a broad range of impact magnitudes, allowing derivation of reliable attenuation factors that were successfully applied to estimate athletes’ striking forces.

### Interpretation of Attenuation Characteristics

The relatively consistent attenuation observed across a wide range of impact magnitudes suggests that the multilayer materials used in Taekwondo body protectors maintained stable energy absorption performance without exhibiting significant material nonlinearity under elevated impact loads. Typical protectors employ multilayer polymeric materials such as EVA foam and polyurethane foam, which are specifically designed to prevent rapid degradation of attenuation performance even after repeated impacts. These material properties likely contributed to the stable attenuation factors observed throughout the repeated impact tests in this study. Furthermore, while the cushioning materials effectively reduced maximum forces, they did not introduce significant dynamic deformation or rebound behavior within the short contact duration, which explains the minimal variation observed in impact duration across conditions.

Previous studies on boxing gloves have reported that repeated impacts lead to convergence of maximum force [38], impact duration, and impulse toward steady-state values. In consideration of this stabilization phenomenon, more than 500 pre-conditioning impacts were applied to the protector prior to attenuation factor measurements in this study.

Another important factor to consider is the fixed condition of the target unit used in the pendulum-based system. In general, when the impacted object remains stationary during collision, higher impact forces are recorded. In actual Taekwondo sparring, however, opponents tend to move backward reflexively upon receiving a strike; to replicate this behavior, the mannequin in the present setup was mounted on a release mechanism that allowed it to pivot and fall backward immediately after contact, thereby permitting additional attenuation through the motion of the target itself. Therefore, the attenuation factors derived in this study may provide a conservative estimate, potentially higher than the forces actually perceived by athletes during live competition.

Material properties and accumulated usage may also influence the attenuation factors depending on the manufacturer or wear history of the protector. While protectors have been introduced to promote objective scoring in Taekwondo, differences in scoring sensitivity and calibration protocols exist across manufacturers. Minimizing these inter-manufacturer discrepancies is desirable for fair competition, and the attenuation quantification methodology proposed in this study could serve as a foundation for establishing standardized performance evaluation criteria for body protectors.

### Interpretation of Load Cell Measurement Results

An adapter cap was mounted on the load cell to expand the sensing area and ensure consistent contact with the protector surface during impact measurements. This configuration demonstrated consistent performance during preliminary manual hammer tests under low-speed impact conditions. However, under high-speed impacts, discrepancies emerged between the impulses measured by the load cell and those estimated from the pendulum’s rebound angle, particularly at higher impact velocities where a plateau in impulse values was observed (Fig 4C). Considering that the bandwidth of the load cell sufficiently covers the frequency range of the impacts applied in this study, these discrepancies are more likely attributed to mechanical factors related to the adapter cap rather than limitations of the sensor’s electronic response. At high impact velocities, minor elastic deformation of the cap or transient micro-gaps between the cap and the load cell sensing surface may act as a mechanical filter, dispersing or attenuating the short-duration maximum forces. Such mechanical effects may have contributed to the observed plateauing of impulse measurements under high-speed conditions. Nonetheless, as both input and transmitted forces were measured using identical load cell configurations with identical adapter caps, the attenuation factors derived from the ratio between these forces are expected to be minimally affected by such artifacts. Therefore, the attenuation factors proposed in this study are considered applicable even under high-speed, high-intensity impact conditions.

The measurement system was designed to accommodate the structural constraints of embedding sensors inside and outside the protector while withstanding repeated high-intensity impacts. This imposed limitations on sensor selection and cap design. Although high repeatability was achieved, factors such as variations in impact angles or asymmetric strikes that may occur in actual competition settings were not fully addressed in this study.

### Validity and Limitations of the Pendulum-Based Impact System

In this study, a free-fall pendulum system was developed to quantitatively analyze the impact attenuation characteristics of Taekwondo body protectors. Compared to conventional impact simulation methods, the system offered substantial improvements in both safety and reproducibility, enabling systematic evaluation of the protector’s attenuation properties and allowing indirect estimation of athletes’ actual striking forces. However, while this system was optimized for deriving attenuation factors, it has limitations in directly evaluating the sensing performance of the embedded electronic scoring systems used in Taekwondo competitions. Accurate evaluation of such scoring sensors requires test systems that more precisely replicate the dynamic characteristics of actual Taekwondo kicks, including not only a range of impact magnitudes but also controlled variations in impact velocity and contact duration.

This study focused on replicating the reported impact-force profiles observed during real Taekwondo kicks. Since the impulse generated by the pendulum is proportional to both its mass and velocity, the metal pendulum—being much heavier than a human foot—would generate excessively high forces if operated at velocities equivalent to real kicks [39]. This would pose substantial challenges to the durability of the target unit. Therefore, the pendulum velocity was intentionally limited to ensure that both maximum forces and impulse values remained within physiologically relevant ranges previously reported for Taekwondo strikes. For the purpose of evaluating scoring system performance, however, it would be necessary to replicate high-velocity, short-duration impacts similar to actual kicks. Achieving such conditions would require a motorized impactor system capable of independently controlling both force magnitude and contact duration, rather than relying on free-fall dynamics. Additionally, as suggested by the plateau phenomenon observed in the high-speed load cell measurements (Fig 4), further development of high-speed impact application and sensing systems is warranted for precise evaluation under such conditions.

### Interpretation of Athlete Striking Force Results

The actual striking forces estimated from the protector’s internal sensor measurements during athlete kicks were approximately 1.5 kN, which falls within the range of impact forces previously reported for Taekwondo kicks. This consistency supports the validity of the attenuation factor-based estimation approach proposed in this study. Unlike the artificial impact tests using mechanical devices, the athlete kicking trials exhibited a distinct double-peak pattern in the force profiles (Fig 7). Similar double-peak phenomena have been reported in previous studies involving impact apparatus with integrated cushioning materials [20, 22, 34]. Two primary mechanisms may explain this observation. First, stress wave reflections may occur when impact-induced stress waves propagate through the compliant backing structures (mannequin and protector) and are reflected back to the load cell with a time delay due to the system’s relatively low stiffness. Second, re-impact or rebound may arise as the elastic or viscoelastic deformation of the compliant mannequin and protector recovers following the initial impact, leading to secondary contact forces. An important consideration is whether the frequency range associated with these double-peak phenomena falls outside the relevant dynamic range of the attenuation factors derived in this study. Given that the cushioning materials generally exhibit limited dynamic frequency-dependent behavior, it is unlikely that these high-frequency components would meaningfully affect the validity of the attenuation factors. Therefore, the attenuation factors proposed here are considered applicable for estimating striking forces during actual athlete kicks despite the presence of double-peak patterns.

## Conclusion

This study systematically quantified the impact attenuation performance of Taekwondo body protectors under repeated dynamic impact conditions and proposed a novel methodology for indirectly estimating athletes’ actual striking forces. The attenuation factor-based estimation approach developed in this study offers a new solution for quantifying striking forces that are otherwise difficult to measure directly, and may serve as a useful tool for performance assessment and training evaluation in Taekwondo athletes. Furthermore, the measurement and analysis framework established here may be applicable to evaluating the impact characteristics of other protective equipment with similar material compositions and structural designs. Future studies may also explore the development of motor-driven impact apparatus capable of replicating a broader range of impact conditions to further advance measurement capabilities.

## Acknowledgments

This work was conducted in collaboration with Korea National Sport University and was supported by a grant from the National Research Foundation of Korea (NRF).

